# Immune Digital Twin Blueprint: A Comprehensive Mechanistic Model of the Human Immune System

**DOI:** 10.1101/2020.03.11.988238

**Authors:** Rada Amin, Sara Sadat Aghamiri, Bhanwar Lal Puniya, Lauren Mayo, Dennis Startsev, Kashish Poore, Resa Helikar, Tomáš Helikar

## Abstract

The immune system is a complex and dynamic network, crucial for combating infections and maintaining health. Developing a comprehensive digital twin of the immune system requires incorporating essential cellular components and their interactions. This study presents the first blueprint for an immune system digital twin, consisting of a comprehensive and simulatable mechanistic model. It integrates 51 innate and adaptive immune cells, 37 secretory factors, and 11 disease conditions, providing the foundation for developing a multi-scale model. The cellular-level model demonstrates its potential in characterizing immune responses to various single and combinatorial disease conditions. By making the model available in easy-to-use formats directly in the Cell Collective platform, the community can easily and further expand it. This blueprint represents a significant step towards developing general-purpose immune digital twins, with far-reaching implications for the future of digital twin technology in life sciences and healthcare, advancing patient care, and accelerating precision medicine.

## Introduction

Digital twin technology has emerged as a powerful tool for creating virtual representations of real-world systems, allowing for the simulation, analysis, and optimization of these systems in a controlled environment. By generating a digital replica of a physical asset, process, or system, digital twin technology enables engineers, scientists, and decision-makers to anticipate problems, design innovative solutions, and assess the viability of new products before their physical implementation^1,2^. The use of digital twins has grown exponentially across various industries, including manufacturing, automotive, and medical devices, due to their potential to improve efficiency, reduce costs, and minimize risks associated with changes to existing operations or the development of new products^3–5^. As real-world data continually informs the digital twin, the accuracy with which it represents the actual system increases, allowing for more informed decision-making and better predictions of system behavior.

Much of the massive influx of data produced by high-throughput technologies in the biomedical domain consists of discrete snapshots of biological processes, offering an incomplete view of working systems. The life sciences will thus benefit greatly from digital twins that can integrate diverse data sources and reconstruct comprehensive, dynamic models of biological systems to facilitate drug discovery, optimize treatment plans, and even replace traditional clinical trials with simulations on virtual patients. Digital twins provide researchers and healthcare professionals with a deeper understanding of the functions and dysfunctions of complex biological systems, ultimately leading to improved patient outcomes and precision medicine^6,7^.

Digital twins have been designed to simulate the progression of individual tumors and personalized cancer scenarios, incorporating parameters like magnetic resonance imaging data, the intricacies of the tumor microenvironment, genetic alterations, a spectrum of omics data, and responses to treatments^8–10^. These models can be used to predict tumor progression, optimize treatment strategies, and identify potential therapeutic targets^11–14^. Digital twins simulating the function of the human heart and its response to various interventions, such as pacemaker settings or drug therapies, and patient-specific digital twins of vascular systems simulating blood flow, pressure, and other hemodynamic parameters can be used to optimize treatment strategies and predict patient outcomes in heart failure, arrhythmias, and other cardiovascular conditions^15–17^. Digital twins have also been developed for patients with type 1 and type 2 diabetes to simulate glucose metabolism, insulin sensitivity, and the effects of various interventions, such as insulin administration and lifestyle modifications, thus helping optimize glucose control and personalize diabetes management strategies^18–21^.

The human immune system is an ideal candidate for digital twin development due to its broad medical significance and complexity^22^. Spanning every level of biological organization, the immune system plays a crucial role in many health conditions, including autoimmune diseases, primary immune disorders, allergies, infections, and systemic biological responses related to chronic diseases, wound healing, and trauma. The immune system is a highly complex and dynamic network involving many cellular and molecular components, such as immune cells, cytokines, and immunoglobulins, which interact in a tightly regulated manner. An accurate understanding of the system requires integrating these components and their interactions into a coherent and consistent model.

This manuscript introduces a comprehensive map and a simulatable (logical) model of the immune system as the first blueprint of an immune system digital twin - a key initial step towards the development of an immune digital twin recently identified by the community^22^:

### Establishing a foundation

The immune system spans different temporal scales and levels of biological organization, including molecular, cellular, tissue, organ, and organism levels. The blueprint provides a solid foundation for a multi-scale model, ensuring that the essential cellular components and their interactions are accurately represented before introducing other levels of biological organization^22^.

### Guiding model development

The blueprint serves as a guide for constructing the multi-scale model, helping researchers identify critical components and interactions that need to be integrated across different scales and ensuring that the model remains consistent and coherent as it expands to encompass additional levels of biological complexity^22^.

### Facilitating validation and refinement

Having a well-defined blueprint allows the community to validate and refine the model more effectively, comparing its predictions with experimental data at different levels of biological organization and making necessary adjustments to improve its accuracy and reliability.

### Enhancing collaboration and interdisciplinary research

A comprehensive blueprint of the immune system can facilitate collaboration among researchers from different disciplines, providing a common framework and language to understand the immune system’s complexity and interactions.

## Methods

### Model construction and mathematical framework

We chose logical modeling to represent the cellular-level interactions within the immune system accurately in the absence of comprehensive quantitative kinetic information. Logical models use rules to describe the relationships between various components of a biological system, such as activation, inhibition, and feedback loops^23–26^. We represented each immune cell type, cytokine, immunoglobulin, and disease as a distinct component, with edges illustrating their interactions. The components are assigned discrete values (e.g., 0 for inactive and 1 for active) based on the presence or absence of a specific component’s activity, while logical rules dictate the state transitions between these values. We built and curated the model in the web-based modeling and analysis platform, Cell Collective^27^. All components and individual interactions used to construct the regulatory mechanisms have been annotated in Cell Collective with the exact quote from the reference literature. A total of 449 scientific publications were used to build the model and have been listed in the reference panel of the Cell Collective platform overview tab. The model is publicly available in Cell Collective, where it can be simulated, further expanded, and downloaded in several file formats (such as SBML-qual ^28^, text files containing the logical functions, and truth tables).

### Model simulations and analyses

We used Cell Collective for all computational simulations and analyses. This platform utilizes discrete mathematics to construct the model, while the simulated output values exhibit semi-continuous behavior, spanning from 0 to 100% activity levels ^25,27^. External components’ activity levels are dimensionless and represent a percentage chance that a component is active at a specific time (*t*). It is important to note that the activity levels provide a semi-quantitative measure of the relative activity of a particular component rather than a specific biological measurement (e.g., concentration). Users can tailor external components’ activity levels as required by the simulation experiment, either by setting specific values or by defining a range from which values are randomly sampled before each simulation (e.g., for simulating dose-response experiments).

Simulations and analyses used asynchronous updates^29,30^. We conducted two types of analyses: real-time and dose-response.

For real-time simulations, the activity of components at different times (steps) was presented using the mean activity level of multiple simulations (mean±standard error of the mean [SEM]). For dose-response experiments, each experiment simulating a single infection consisted of 100 simulations with different randomly sampled external component activity levels. Each simulation consisted of 5,000 steps. Output components’ activity levels were calculated as the fraction of 0’s and 1’s over the last 500 iterations^25,31,32^. As noted previously^25,32^ and from observations drawn from the model presented here, the model reaches a steady state rapidly, and these values are sufficient to describe the network’s “long-term” (e.g., attractor-like) behavior.

### Simulation settings

Each pathogen is represented by an external component (independent variable) in the model whose activity level can be set by the user. Each immune cell also possesses an associated external component so that its initial levels can be set to represent different immune system health statuses for various conditions or sub-populations. For real-time simulations on the Cell Collective platform, under the “Simulation” tab, the “External Components”, the simulations have been set to 100% to simulate the presence of specific external component(s) activity level. For dose-response analysis, the model was simulated under 67-100% pathogens’ activity levels. The simulations we defined mimic real-world scenarios, categorizing the severity of infections into different stages (stage 1 (1-34%), stage 2 (34-67%), and stage 3 (67-100%)). The range used for the simulations is based on the highest pathogen load and the impacts on host physiology^33,34^.

### Statistical analysis

Statistical analyses were performed with GraphPad Prism software using the unpaired parametric Student two-tailed *t-test* as appropriate.

### Code availability

The model is freely available on the Cell Collective platform (contact authors for direct link) This platform is a user-friendly online environment for building, simulating, and analyzing computational models of biological systems, allowing researchers to access, modify, and utilize the model for their research questions.

## Results

### Model design and scope

Our comprehensive model captures the intricate network of signals and responses that regulate the immune system’s defense against disease conditions and comprises 124 components representing: disease/pathogen target cells; innate and adaptive cells and their respective subtypes (e.g., T helper [Th] 1, 2, 9, 17, 22 for CD4 T cells) and their various states (e.g., resting, naive, activated, antigen presentation); 37 secretory factors such as interleukins (ILs), immunoglobulins (Igs), growth factors, and reactive oxygen species (ROS); 9 pathogens; an autoimmune disease (type 1 diabetes, T1D); transplant (lung transplantation, LTx); and 1,450 regulatory interactions among these components (Fig. 1a). A schematic overview of the model components, with the number of subtypes per component stated in parentheses, is provided (Fig. 1b), along with a detailed description of each cell type and subtype included in the model, their definition, and associated references (Table 1).

**Figure 1:**
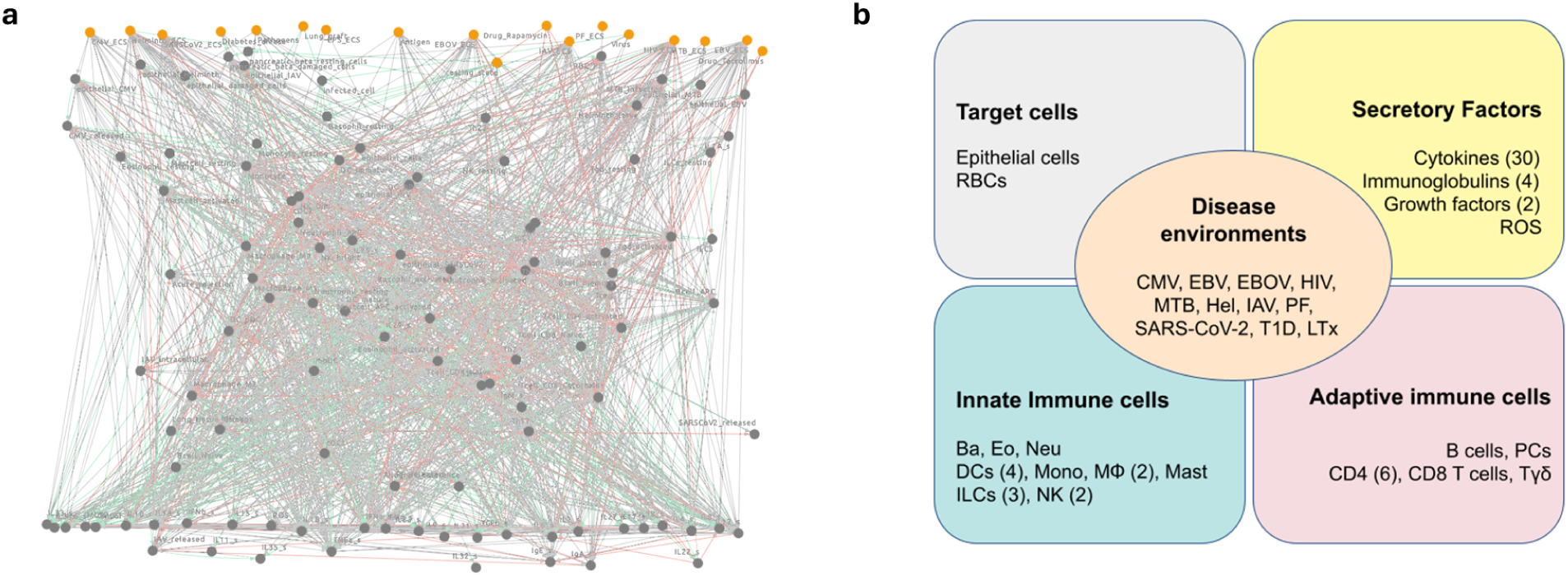
Components of the immune system model. **a** Network visualization of digital twin blueprint of the human immune system in Cell Collective. **b** Overview of the cell-cell model components, with the number of subtypes per cell type mentioned in parentheses. Disease environments: Cytomegalovirus (CMV), Epstein-Barr Virus (EBV), Ebola virus (EBOV), Human Immunodeficiency Virus (HIV), Mycobacterium tuberculosis (MTB), Helminth (Hel), Influenza A virus (IAV), Plasmodium Falcipurum (PF), Severe acute respiratory syndrome coronavirus 2 (SARS-CoV-2), Type 1 Diabetes (T1D), Lung transplantation (LTx). Secretory factors: Reactive oxygen species (ROS). Target cells: Red blood cells (RBCs). Innate cells: Basophils (Ba), Eosinophils (Eo), Neutrophils (Neu), Dendritic cells (DCs), Monocytes (Mono), Macrophages (MՓ), Neutrophils (Neu), Mast cells (Mast), Innate lymphoid cells (ILCs), Natural Killer cells (NK). Adaptive cells: Plasma cells (PCs), gamma-delta T cells (Tγδ).

**Table 1:** List of cell types included in the modeled immune system.

| Cell Types | Brief Synopsis | Citations |
| --- | --- | --- |
| **Target Cells** |  |  |
| Epithelial cells | These barrier-forming cells are essential for maintaining tissue integrity and preventing pathogen entry. They secrete interleukins, chemokines, and growth factors, such as IL-1, IL-6, IL-8, TGF-β (transforming growth factor-β), and GM-CSF (granulocyte-macrophage colony-stimulating factor), that recruit and activate immune cells, such as neutrophils, macrophages, and dendritic cells. | 96 |
| Red blood cells (RBCs) | These cells transport oxygen, carbon dioxide, and nutrients throughout the entire body. Modifications in their structure and quantity serve as clinical indications of disease processes. | 97 |
| **Innate Immunity** |  |  |
| Basophils | These granulocytes contribute to allergic reactions and inflammation. They are activated when antigens cross-link IgE bound to their FcεRI receptors. Upon activation, basophils release histamine and other inflammatory mediators, including IL-4, IL-13, and IL-33, that increase vascular permeability and attract other immune cells, such as eosinophils, to the site | 98,99 |
|  | of inflammation. |  |
| Dendritic cells (DCs) | These antigen-presenting cells (APCs) serve as a crucial bridge between innate and adaptive immunity. Their primary function involves capturing and processing antigens, which are then presented to T cells through major histocompatibility complex (MHC) molecules, initiating adaptive immune responses. Activation of DCs occurs upon recognition of pathogen-associated molecular patterns (PAMPs) or damage-associated molecular patterns (DAMPs) through pattern recognition receptors (PRRs) like toll-like receptors. Subsequently, DCs release a variety of cytokines, such as IL-12, IL-6, and IL-1, which promote T cell differentiation and activation, thus enhancing the adaptive immune response. DCs are divided into three major subtypes: monocyte-derived DCs (moDCs), conventional DCs (cDC1 and cDC2), and plasmacytoid DCs (pDCs). | 30,100,101 |
| Monocyte-derived dendritic cells (moDCs) | These DCs are derived from monocytes in response to inflammation and infection. They activate both CD8+ and CD4+ T cells through antigen presentation and the secretion of TNF-α, IL-1β, and IL-12, which are essential for driving inflammatory responses and promoting T cell polarization. | 40,102 |
| Conventional type 1 dendritic cells (cDC1) | cDC1 cells are crucial in initiating anti-viral immunity by efficiently cross-presenting antigens to CD4+ and CD8+ T cells. They produce high levels of IL-12, which is important for driving Th1 responses and enhancing the function of cytotoxic T cells in killing infected cells. | 39,40 |
| Conventional type 2 dendritic cells (cDC2) | These subtypes of DCs initiate immune responses against extracellular pathogens, such as bacteria and fungi. They produce various cytokines, including IL-6, IL-12, IL-23, and IL-10, which help shape the immune response by promoting CD8+ T cell, Th2, Th17, and regulatory T cell responses. | 39,40 |
| Plasmacytoid dendritic cells (pDC) | These DCs primarily sense viral infections through toll-like receptors (TLRs), particularly TLR7 and TLR9, which recognize viral RNA and DNA, respectively. Upon viral infection, pDCs rapidly produce large amounts of IFN-γ, which is responsible for the activation and proliferation of NK cells, T cells, and cDCs. | 103,104 |
| Eosinophils | These granulocytes defend against parasitic infections and contribute to allergic inflammation. Eosinophils are activated by cytokines produced by Th2 cells, such as IL-5, and by other inflammatory mediators, such as leukotrienes. Upon activation, they release cytotoxic granules containing reactive oxygen species (ROS) and release cytokines, such as IL-4, that modulate other immune cell functions. | 105 |
| Innate lymphoid cells (ILCs) | These cells defend against diverse infections and play a role in initiating adaptive responses. Presented as an innate counterpart of CD4+ T helper cells, ILCs share a similar array of cytokines upon activation by epithelial- or myeloid cell-derived cytokines. They promote inflammation, mucus production, and tissue repair. ILCs comprise three subtypes: ILC1, ILC2 and ILC3. | 106 |
| Type 1 innate lymphoid cells (ILC1s) | These cells are crucial in early defense against viral infection as they can directly kill infected cells through the production of cytotoxic mediators, including granzymes and perforins. ILC1s secrete Th1 cytokines that mirror those produced by CD4+ T cells, including the production of IFN-γ and TNF-α. | 107 |
| Type 2 innate lymphoid cells (ILC2s) | ILC2s are the innate counterparts of Th2 cells and protect against parasite infections. They secrete cytokines similar to Th2 cells, including IL-4, IL-13, and IL-5, that promote the recruitment of eosinophils and mast cells and facilitate the differentiation of M2 macrophages and plasma cells. | 108 |
| Type 3 innate lymphoid cells (ILC3s) | Presented as sentinel cells in mucosa, these cells mainly defend against bacteria and fungi. By secreting similar cytokines to Th17 cells, including IL17, IL-22, and GM-CSF, ILC3s promote neutrophil recruitment, plasma cell differentiation, and myeloid activation. | 109 |
| Macrophages | These phagocytic cells engulf and clear pathogens, remove cellular debris, and produce cytokines that regulate inflammation and immune responses. Macrophages can be activated by PAMPs, DAMPs, or cytokines, such as IFN-γ (interferon-γ) and GM-CSF. Upon activation, macrophages release a diverse array of cytokines including IL-1, IL-6, TNF-α (tumor necrosis factor-α), IL-10, and IL-12, as well as other factors like nitric oxide, ROS, and inducible nitric oxide synthase (iNOS). These molecules collectively modulate immune cell activation and differentiation, contributing to the orchestration of immune responses. Macrophages comprise two main subtypes: M1 and M2 macrophages. | 110 |
| M1 macrophages | M1 macrophages are classified as pro-inflammatory cells and produce a variety of pro-inflammatory cytokines, such as IL-1, IL-6, IL-12, and TNF-α, upon microbial infection or by the presence of IFN-γ in the milieu. These cytokines create an inflammatory milieu that enhances the recruitment and activation of other immune cells, such as T cells and NK cells. Additionally, the high levels of reactive oxygen species (ROS) and nitric oxide (NO) produced by M1 macrophages contribute to pathogen clearance and the immune response. | 111 |
| M2 macrophages | In contrast to M1 macrophages, M2 macrophages are anti-inflammatory cells that resolve inflammation through the production of anti-inflammatory cytokines. The cytokines produced by M2 macrophages, including IL-4, IL-13, IL-10, and TGF-β, foster an anti-inflammatory environment that inhibits the activity of pro-inflammatory cells and promotes the resolution of inflammation. M2 macrophages are essential in tissue repair, homeostasis, and wound healing after an inflammatory response. | 111 |
| Mast cells | These cells play a role in allergic reactions and inflammation. Upon crosslinking of the IgE-antigen complex to the surface receptor FcεRI, mast cells release histamine, tryptase, and other mediators. Mast cells also produce cytokines, such as TNF-α, IL-4, IL-5, IL-6, and IL-13, which modulate the function of other immune cells, including T cells and DCs. | 112 |
| Monocytes | These circulating cells differentiate into macrophages or DCs upon entering tissues. They participate in phagocytosis and release cytokines, such as IL-1, IL-6, TNF-α, and IL-10, which regulate inflammation and immune responses. Monocytes can be activated by PAMPs, DAMPs, or cytokines, such as GM-CSF and M-CSF (macrophage colony-stimulating factor). | 113 |
| Neutrophils | These phagocytic cells are the first responders to infection and tissue damage. They release antimicrobial peptides, proteases, and ROS to destroy pathogens. Neutrophils are rapidly recruited to sites of infection or inflammation by chemokines, such as IL-8 and MIP-1α, and other inflammatory mediators produced by epithelial cells, endothelial cells, and other immune cells. | 114 |
| Natural Killer (NK) cells | These cytotoxic cells recognize and directly kill virus-infected and cancerous cells without prior sensitization. Activation of NK cells occurs through various means, including cytokine stimulation, such as IL-12, IL-15, and IL-18, as well as recognition of stress-induced ligands present on target cells. Upon activation, NK cells not only perform cytotoxic functions but also secrete immunomodulatory cytokines like IFN-γ. The cytokine production by NK cells promotes Th1 responses and augments the cytotoxic activity of CD8+ T cells. | 115,116 |
| NK bright cells | These cells demonstrate an immunomodulatory role by secreting both pro- and anti-inflammatory cytokines, including IFN-γ, TNF-α, IL-10, and GM-CSF. NK bright cells are described as immature and with low cytotoxic ability, but upon infection they can differentiate into a mature NK dim cell that displays higher cytotoxic potential. | 47,48 |
| NK dim cells | These cells can directly induce apoptosis of infected cells through the release of cytotoxic granules and initiation of antibody-dependent cellular toxicity (ADCC). NK dim cells can also secrete IFN-γ and TNF-α but to a lesser extent than NK bright cells. These cytokines play significant roles in promoting immune cell recruitment and enhancing antigen presentation capabilities of DCs and macrophages. | 48 |
| **Adaptive Immunity** |  |  |
| B cells | B cells are integral to the adaptive immune system, expressing membrane-bound antibodies called immunoglobulins (Ig) (such as IgM, IgA, IgE, and IgG classes) that target and neutralize specific pathogens. Additionally, activated B cells can serve as antigen-presenting cells for T cells, further enhancing the coordination of immune responses. | 117 |
| Plasma cells (PCs) | Upon encountering their specific antigen, B cells undergo differentiation into antibody-secreting PCs, which is mediated by CD4+ T cells. This immunological synapse relies on the interaction between CD40 ligand (CD40L) on T cells and CD40 on B cells, complemented by the release of cytokines, such as IL-4, IL-6, and IL-21. | 117 |
| CD4+ T cells | Upon activation by DCs presenting antigens via MHC II, naive CD4+ T cells can differentiate into various Th cell subsets, such as Th1, Th2, Th9, Th17, Th22, and regulatory T cells (Tregs). This differentiation is influenced by the cytokine milieu produced by other immune cells, such as DCs, | 118,119 |
|  | macrophages, and NK cells. |  |
| T helper 1 (Th1) cells | These CD4+ T cells are involved in cell-mediated immunity against intracellular pathogens, such as viruses and some bacteria. They produce IFN-γ and TNF-α, which promote macrophage activation and enhance the cytotoxic activity of CD8+ T cells and NK cells. Th1 cell differentiation is induced by IL-12 and IFN-γ that are released by antigen-presenting cells, such as DCs and macrophages, in response to intracellular pathogens. | 92,118 |
| T helper 2 (Th2) cells | These CD4+ T cells are involved in humoral immunity against extracellular pathogens, such as parasites. They produce IL-4, IL-5, and IL-13, which promote B cell differentiation into plasma cells, eosinophil activation, and mast cell activation. Th2 cell differentiation is induced by IL-4, which can be released by DCs, basophils, or mast cells. | 118,120 |
| T helper 9 (Th9) cells | These CD4 + T cells are characterized by their ability to produce IL-9, a cytokine involved in various immune responses. Th9 cells play a role in immune-related diseases such as inflammation, allergy, and parasite infection. | 121 |
| T helper 17 (Th17) cells | These CD4+ T cells play a role in defending against extracellular bacterial and fungal infections. They produce cytokines, such as IL-17A, IL-17F, IL-21, and IL-22, which stimulate the production of antimicrobial peptides and the recruitment of neutrophils. Th17 cell differentiation is induced by the cytokines IL-6, TGF-β, IL-1, and IL-23. | 120,122 |
| T helper 22 (Th22) cells | These cells are named after the main cytokine that they produce, IL-22. These CD4+ T cells are involved in immune responses at barrier surfaces, such as the skin and mucosal membranes, and contribute to defense against certain infections including HIV and influenza. Additionally, Th22 cells have been implicated in various inflammatory and autoimmune conditions. | 123 |
| Regulatory T cells (Treg) | These CD4+ T cells play a critical role in maintaining immune tolerance and preventing autoimmunity. They suppress the activation and function of other immune cells, including B cells, CD4+ T cells, CD8+ T cells, and DCs, through various mechanisms, such as the secretion of immunosuppressive cytokines (e.g., TGF-β and IL-10) and direct cell-cell interactions. Treg differentiation is induced by TGF-β and IL-2. | 120,124 |
| CD8+ cytotoxic T cells | These cytotoxic T cells recognize and kill virus-infected and cancerous cells by releasing cytotoxic granules containing perforin and granzymes. CD8+ cytotoxic T cell activation requires antigen presentation by MHC I molecules on infected or abnormal cells, as well as help from CD4+ T cells in the form of cytokines, such as IL-2 and IFN-γ. | 37,125 |
| Gamma-delta ($\gamma\delta$) T cells | These cells represented a minor subset of T cells, with the capability to present antigens similar to innate cells. These cells present dual functions in maintaining immunosuppression or activating inflammation due to the ambivalent cytokine function they produce, including IL-4, IL-17, IL-21, IL-22, GM-CSF, and IFN-γ. | 126 |

### Immune system stimuli

As stated, the model encompasses an autoimmune disease (i.e., T1D), LTx, and nine common pathogens from different viral, bacterial, and parasitic groups to simulate the initiation of the immune response under various disease conditions (Table 2). By incorporating pathogen-specific immune responses and host-pathogen interactions at the cellular level, the model provides a realistic representation of the complex interplay between the immune system and invading pathogens.

**Table 2:** List of disease conditions of the modeled immune system.

| Disease Environments | Brief Synopsis | Citations |
| --- | --- | --- |
| Cytomegalovirus (CMV) | Cytomegalovirus (CMV) triggers an innate response involving DCs and NK cells, leading to the release of type I interferons to regulate the early stages of infection. During the persistence phase, CD4+ T cells, CD8+ T cells, and B cells, primed by NK cells and DCs, orchestrate a vigorous and enduring immune response to restrain high viral loads. Despite this response, CMV can establish a latent state within the host, posing a risk of reactivation and reinfection particularly when the immune system is compromised. | 127 |
| Epstein-Barr virus (EBV) | Epstein-Barr virus (EBV) primarily infects B cells, which can be eliminated by NK cells, effector CD4+ T cells, and CD8+ T cells, thereby controlling the initial phase of infection and maintaining viral latency. However, sporadic reactivation may occur, evading immune surveillance despite the presence of long-lasting CD8+ and CD4+ T cell responses. | 128,129 |
| Ebola virus (EBOV) | Ebola virus (EBOV) triggers a multifaceted immune response, where innate defenses mobilize macrophages, DCs, and NK cells to release proinflammatory cytokines such as TNF-α, IL-6, and IFN-$\alpha/\beta$. In the adaptive arm, CD8+ T cells directly target infected cells, while CD4+ T cells secrete key cytokines like IFN-γ and IL-2. Additionally, plasma cells produce neutralizing antibodies to counteract the virus. | 130,131 |
| Human immunodeficiency virus (HIV) | The immune response against Human Immunodeficiency Virus (HIV) comprises both innate and adaptive components. In the innate immune response, DCs, NK cells, and macrophages produce IFN-$\alpha/\beta$ and other cytokines. Adaptive immunity involves CD8+ T cells, which directly target infected cells, and CD4+ T cells, which secrete cytokines such as IFN-γ and IL-2. Plasma cells contribute by producing neutralizing antibodies. However, HIV's high mutation rate enables it to evade the immune system, posing a significant challenge for effective immune responses. | 132 |
| Influenza A virus (IAV) | The immune response against influenza A virus (IAV) commences with the activation of the innate system, comprising macrophages, DCs, and NK cells, which release cytokines including IFN-$\alpha/\beta$, TNF-α, and IL-6. Adaptive immunity then follows, with CD4+ T cells producing cytokines like IFN-γ and IL-2 to orchestrate the immune response, while CD8+ T cells target and eliminate infected cells. Additionally, plasma cells produce neutralizing antibodies that specifically recognize and bind to viral surface proteins, such as hemagglutinin and neuraminidase, aiding in viral clearance. | 133 |
| Mycobacterium tuberculosis (MTB) | The immune response to mycobacterium tuberculosis (MTB) entails macrophages engulfing the bacteria and producing cytokines like TNF-α, IL-1, and IL-6. CD4+ T cells produce IFN-γ and IL-2, activating macrophages to eliminate intracellular bacteria. Simultaneously, CD8+ T cells target and eradicate infected cells. Although plasma cells produce antibodies, their precise role in MTB immunity remains incompletely | 134,135 |
|  | understood. |  |
| Severe acute respiratory syndrome coronavirus 2 (SARS-CoV-2) | COVID-19, caused by the SARS-CoV-2 virus, induces the activation of neutrophils, monocytes, and macrophages. In severe cases of COVID-19, inflammation disrupts both innate and adaptive immune responses, resulting in an excessive release of cytokines such as IL-6, IL-1β, and TNF-α. These cytokines are implicated in the development of clinical complications associated with the disease. | 136,137 |
| Helminth | These parasitic worms trigger a multifaceted immune response. Initially, the innate immune system responds by activating group 2 innate lymphoid cells (ILC2) and eosinophils, which release a cascade of cytokines including IL-4, IL-5, IL-9, and IL-13. These cytokines play crucial roles in orchestrating defense against helminth infections by promoting mucus production, smooth muscle contraction, and recruitment of additional immune cells to the site of infection. Subsequently, the adaptive immune system mounts a Th2 response, characterized by CD4+ T cells secreting similar cytokines as those produced by the innate immune cells. | 138,139 |
| Plasmodium falciparum (PF) | The pathogen plasmodium falciparum (PF), the causative agent of severe malaria, elicits a complex immune response. In the innate arm, macrophages and neutrophils engage in phagocytosis of infected RBCs, attempting to curb parasite proliferation. DCs and NK cells contribute by releasing key cytokines such as IFN-γ and TNF-α, which aid in parasite clearance. The adaptive immune response is orchestrated by CD4+ T cells, which produce a spectrum of cytokines including IFN-γ, IL-2, and IL-10, crucial for both parasite control and regulation of inflammation. Additionally, plasma cells play a pivotal role by generating antibodies specifically targeting the parasite, thus assisting in the immune-mediated elimination of PF. | 140,141 |
| Type 1 diabetes (T1D) | Type 1 diabetes (T1D) is characterized by an autoimmune attack against the insulin-producing beta cells in the pancreas. The immune response in T1D involves the activation of both CD4+ and CD8+ T cells, which erroneously identify beta cell antigens as foreign and mount an inflammatory response against them. This inflammatory cascade is further fueled by the secretion of cytokines such as IL-6, IL-17, and IL-21. Additionally, autoantibodies produced by plasma cells target beta-cell antigens, contributing to the ongoing destruction of pancreatic beta-cells. This immune-mediated destruction ultimately leads to insulin deficiency and the clinical manifestations of T1D. | 142 |
| Lung transplantation (LTx) | As a result of solid organ transplantation, the recipient's immune system can perceive the newly transplanted lung as foreign tissue, initiating an immune response. This response engages various immune cells, including T cells, B cells, and antigen-presenting cells, which collaborate to mount an attack against the transplanted lung tissue. Pro-inflammatory cytokines produced during this process exacerbate inflammation, increasing the risk of rejection of the transplanted lung. Managing this immune response and preventing rejection are critical aspects of | 143 |
|  | post-transplant care in LTx. |  |

These pathogens and disease conditions prompt a dynamic array of immune responses, engaging both the innate and adaptive arms of the immune system. Innate immune cells, such as macrophages, dendritic cells (DCs), neutrophils, and natural killer (NK) cells swiftly recognize and respond to invading pathogens through mechanisms like phagocytosis, cytokine secretion, and direct cytotoxicity. Simultaneously, adaptive immune cells, including T lymphocytes (CD4+ and CD8+ T cells) and B lymphocytes, undergo activation and differentiation to mount targeted and specific responses against the pathogen. This immune activation triggers the release of a myriad of signaling molecules, including cytokines, chemokines, and other soluble mediators, which orchestrate the immune response. Cytokines such as interleukins (ILs), interferons (IFNs), and tumor necrosis factor (TNF) regulate inflammation, cell proliferation, and differentiation, while chemokines guide immune cell migration to sites of infection.

Understanding the intricacies of immune responses is paramount for the development of effective treatments and vaccines to combat diseases. Insights into how pathogens interact with the immune system, evade immune surveillance, and induce pathology inform the design of targeted therapeutics, including antiviral drugs, immunomodulators, and vaccines. By deciphering the complex interplay between pathogens and the immune system, researchers can devise strategies to enhance host defense mechanisms, mitigate disease progression, and ultimately safeguard global health.

### Model validation: Immune response to pathogens

To evaluate the accuracy and reliability of the computational model, we compared its predictions with published human data, including *in vitro* and *ex vivo* studies and clinical observations. These data encompass various aspects of immune responses to select pathogens making them a suitable blueprint for validating the computational model.

We first evaluated the model’s capacity to replicate inherent cell responses during IAV using real-time simulations. Jost et al. previously found a decline in NK bright cells and an increase in activated NK dim cells among patients infected with seasonal IAV or the H1N1 strain, and our immune system model was constructed based on the experimental data from H1N1 infected patients^35^. We validated the NK cell phenotype during IAV infection by running real-time simulations in our model. Results from the simulation mirror the published findings by predicting a reduction in NK bright cells during acute IAV infection (Fig. 2a), thus demonstrating the accuracy of the model.

**Figure 2:**
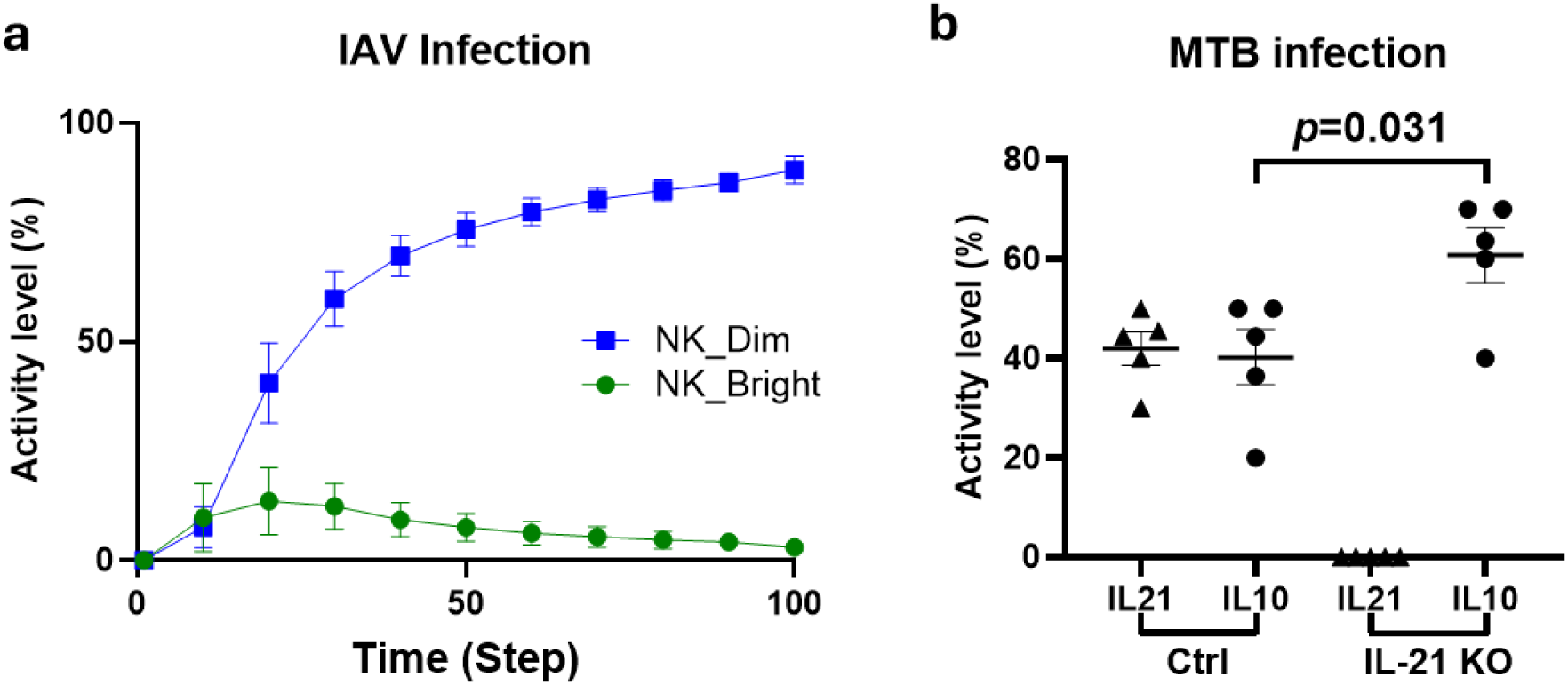
*In silico* validations. **a** Time course distribution of NK bright (green) and NK dim (blue) cells during IAV infection. **b** Assessment of IL-21 (triangle) and IL-10 (circle) activity levels upon MTB infection with control (Ctrl) or without IL-21 (IL-21 KO). (replicates n=5). Data are presented as mean±SEM, and the *p*-value is determined by an unpaired two-tailed t-test.

We further assessed the interaction between cytokines in MTB infection since cytokine activity is an essential regulator of immune response. Previously, Paidipally et al. found that IL-21 siRNA enhanced IL-10 production by peripheral blood mononuclear cells infected with MTB^36^. To address whether this phenomenon could be replicated in our model, we simulated MTB infection and evaluated the activity level of IL-21 and IL-10 in the absence or presence of IL-21 activation. Consistent with the published findings, our model predicted that the absence of IL-21 activity resulted in a significant increase in IL-10 levels during MTB infection (SEM (20.56 ± 7.882). p=0.031) (Fig. 2b). Together, these results demonstrate the model’s ability to accurately simulate immune cell response and cytokine production following exposure to specific pathogens.

Comparing the model’s output with published human findings assesses its ability to appropriately simulate innate and adaptive immune cell dynamics during infection. As shown in Table 3, our model predicted the immune response of multiple innate and adaptive cell subtypes following exposure to select pathogens, which was supported by published literature. Specifically, we presented the reactions of several innate cells, including DCs, macrophages, monocytes, neutrophils, and NK cells, to nine pathogens, and all pathogens demonstrated activation of DCs and/or macrophages. This is attributed to their critical role as antigen-presenting cells (APCs), which is indispensable for initiating the adaptive T cell response. In addition to DCs and macrophages, the simulation showed the activation of neutrophils, NK cells, and monocytes in most pathogens, alluding to their critical role in pathogen clearance through phagocytosis and cytokine release (Tables 1 and 2). For the adaptive response, both CD8+ T cells and the humoral response are activated under all pathogen conditions. The cytotoxic function of CD8+ T cells is essential in host defense against pathogens through the elimination of infected cells, whereas the antibody response plays a crucial role in pathogen opsonization for phagocytosis and antibody-dependent cellular toxicity (ADCC)^37^. Most pathogens displayed CD4+ Th1-specific immune responses, whereas helminths showed CD4+ Th2-specific immune responses. HIV did not initiate CD4+ Th1- or Th2-specific immune responses, and this low activation of immune cells (e.g., CD4+ T cells) mimics the stage of disease with immunodeficiency. The model appropriately predicted that the Th1 response is primarily triggered by bacterial and viral infections, whereas Th2 is activated in the presence of parasitic infections^38^.

**Table 3:** Innate and adaptive cell response to nine different pathogens at the single scale infection as predicted by our model.

| **Infectious Agents Modeled** | **Innate Immune Responses Predicted** | **Adaptive Immune Responses Predicted** | **Supporting References** |
| --- | --- | --- | --- |
| CMV | DCs, NK cells | Th1, CD8+, IgM, IgA, IgG | 43,144,145 |
| EBV | DCs, Monocytes, Macrophages, NK cells, Neutrophils | Th1, CD8+, IgM, IgA, IgG | 146,147 |
| EBOV | DCs, Macrophages, Monocytes | Th1, CD8+, IgM, IgA, IgG | 148 |
| HIV | DCs, Macrophages | CD8+, IgM, IgA, IgG | 149 |
| IAV | DCs, NK cells | Th1, CD8+, IgM, IgA, IgG | 150 |
| SARS-CoV-2 | DCs, Neutrophils, Monocytes, Macrophages | CD8+, IgM, IgA, IgG | 136,137 |
| MTB | DCs, Macrophages, Neutrophils | Th1, CD8+, IgM, IgA, IgG | 151 |
| Helminth | DCs, Macrophages, | Th2, IgM, IgA | 53 |
| PF | DCs, Monocytes, Macrophages | Th1, IgM, IgG, IgA | 152,153 |

Next, we simulated cytokine and Ig production in response to each infection (Fig. 3) and compared our results with the literature. CMV infection induces a stronger inflammatory cytokine response (IL-1ß, IL-2, IL-6, IL-12, IL-15, IFN-γ, TNF-α, and TGF-ß) with low expression of IgE in our model, which is aligned with the literature^39,40^. EBV infection stimulates granulocyte-macrophage colony-stimulating factor (GM-CSF) and several pro-inflammatory cytokines, including IL1-ß, IL-6, IL-8, IL-18, TNF-α, IFN-α, IFN-ß, and IFN-γ, with low levels of IL-4. This response predicted by our model is also aligned with experimental data^41,42^. Our model predicted that EBOV infection promotes a strong response of both pro-inflammatory cytokines (IL-1ß, IL-6, IL-12, IL-15, IFN-α/ß/γ, TNF-α) and anti-inflammatory cytokines (IL-8 and IL-10), which mimics previously published results^43^. HIV infection in our model is characterized by a burst of cytokines, including pro-inflammatory (IL-1α/ß, IL-2, IL-6, TNF-α, IFN-α/ß/γ) and anti-inflammatory (IL-4, IL-10, IL-13) mediators; however, some cytokines did not show any activity level, including IL-9 and IL-22, which were previously shown to be reduced during HIV progression^41,44,45^. IAV infection occurs in alveolar macrophages in the lower respiratory tract and induces a robust pro-inflammatory response (IL-1α/ß, IL-2, IL-5, IL-6, IL-10, IL-12, IL-17, IL-18, IFN-γ, TNF-α, and IL-23)^46^, which was confirmed by our model. Concerning SARS-CoV-2, our simulation exhibits a large panel of cytokines that overlap with those published, including IL-1ß, IL-2, IL-3, IL-4, IL-5, IL-6, IL-8, IL-10, IL-12, IL-17, IL-18, TNF-α, IFN-γ, and GM-CSF^47–49^. This mirrors the cytokine storm and subsequent severe inflammation, immune dysfunction, and tissue damage seen in SARS-CoV-2^47,50^. The model found that MTB elicits a strong pro-inflammatory cytokine response, increasing IL-1α/ß, IL-2, IL-6, TNF-α, IFN-α/ß/γ, and IL-23, and inducing some cytokines with dual functions such as IL-22, IL-27, and IL-35, which also aligns with the literature^51^. Our simulations found that Helminth infection prompts the production of key cytokines, such as IL-4, IL-3, IL-5, and IL-13 (Fig. 3), that are associated with Th2 responses (Table 3)^52^. In contrast to other pathogens, IgG is inactive in helminth infection, consistent with prior research findings that indicate these antibodies are susceptible to enzymatic cleavage as a strategy to evade ADCC^53^. Cytokine response to PF is mixed pro- and anti-inflammatory in the model, which aligns with the literature since both pro- and anti-inflammatory responses are important in the immune response against malaria^54^. Additional cytokine validations are available in Table 1 . Among all pro-inflammatory cytokines, IL-1ß, IL-6, and IFN-γ were identified as the most active in response to simulated infections by different pathogens. Although these cytokines play a protective role against infection, their excessive secretion has been associated with high inflammation, damage, and dysfunction of immune responses in infectious diseases^55–57^. Conclusively, the model simulates key cytokine and Ig activity following pathogen exposure that aligns with current studies.

**Figure 3:**
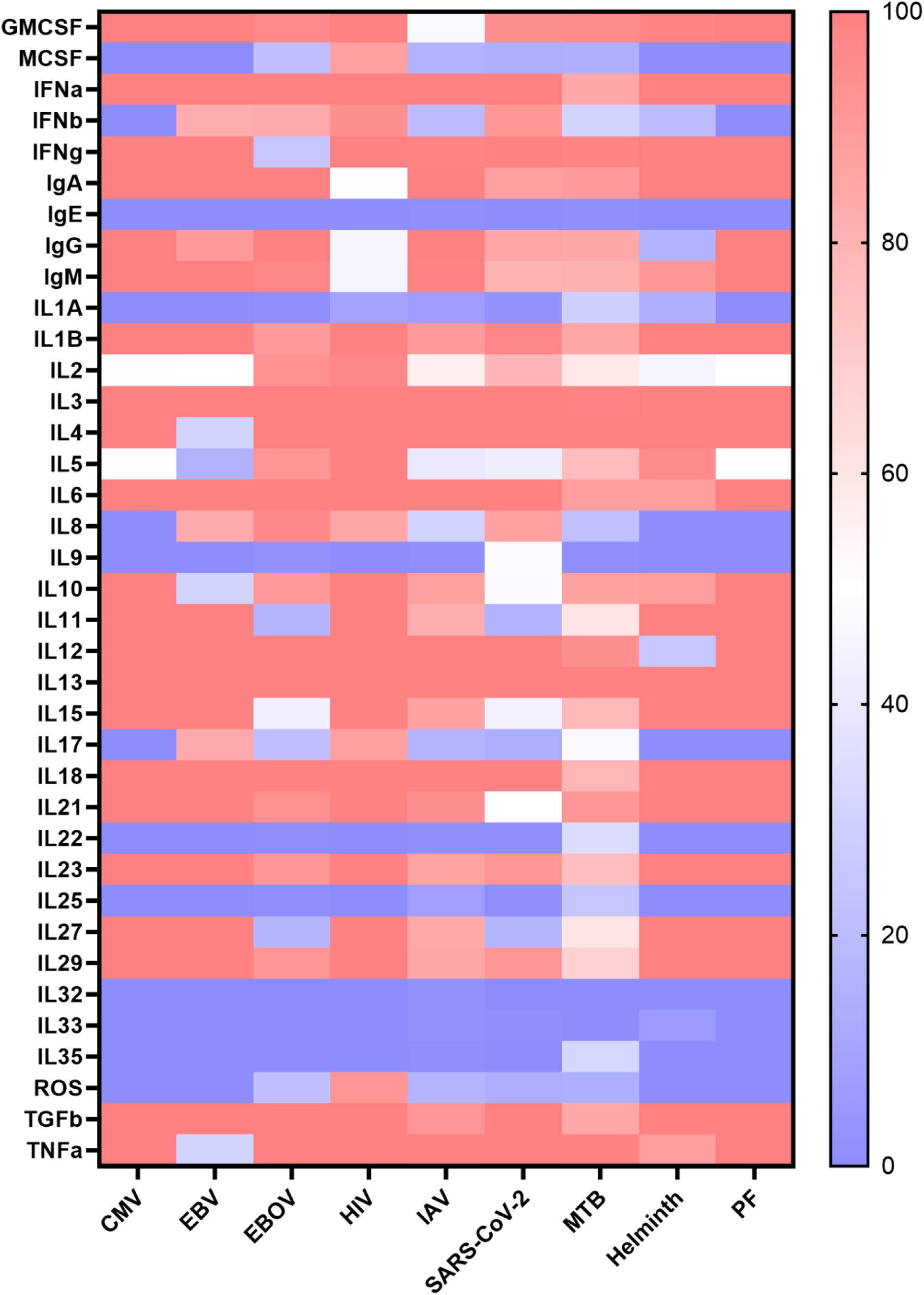
Dose-response analysis of the secretory response to single pathogen infections. (A) Cytokine activity upon infection. The activity levels represent the average value of 100 simulation results triggered by each pathogen and were performed at 67-100% pathogen activity level.

Despite the extensive literature on these nine pathogens, certain cytokines, such as IL-9, IL-32, and IL-35, lack experimental validations that explore the intricate chain linking pathogen-cell-cytokine responses. Consequently, the model’s scope is constrained by the absence of data in the literature concerning any unverified interactions. In summary, the model accurately replicated the immune responses for nine different pathogens, and correctly simulated cytokine and Ig production in response to each infection, aligning with the literature. Notably, the model appropriately reproduced complex experimental scenarios, such as the increase of IL-10 in the absence of IL-21 during replication of MTB infection in the absence of IFN-γ (Fig. 2a). Overall, the model showed a high level of accuracy and reliability in simulating immune responses to infections.

### Case study 1: Immune response to various coinfections

Coinfections can complicate clinical presentation, diagnosis, and treatment, often resulting in more severe symptoms and increased morbidity and/or mortality, underlining the importance of understanding the immune response during coinfection. Characterizing immune responses to coinfections allows researchers to identify key differences in cytokine and Ig activity depending on the infection order leading to more effective therapies and targeted interventions, thereby improving patient outcomes. Furthermore, examining the immune system’s behavior during coinfections provides valuable information on the complex interplay between various pathogens and the host’s immune system. This knowledge can help researchers better understand the mechanisms behind immune system regulation, identify potential vulnerabilities, and design innovative strategies to prevent or manage coinfections. In an era of emerging and re-emerging infectious diseases, understanding the immune responses to coinfections is paramount to global public health.

To address these issues, we analyzed the model’s dynamics in response to medically relevant (observed) coinfections. First, we explored the CD4+ Th1 response in MTB and HIV coinfection, using dose-response analysis, since some clinical observations showed that HIV infection induces the decline of Th1 response in coinfected patients, increasing the susceptibility to MTB infection^58,59^. The model simulation aligned with the literature, confirming the decrease of Th1 activity in HIV-MTB coinfection compared to MTB single infection (SEM (-53.1±0.43), p<0.0001) (Fig. 4a).

**Figure 4:**
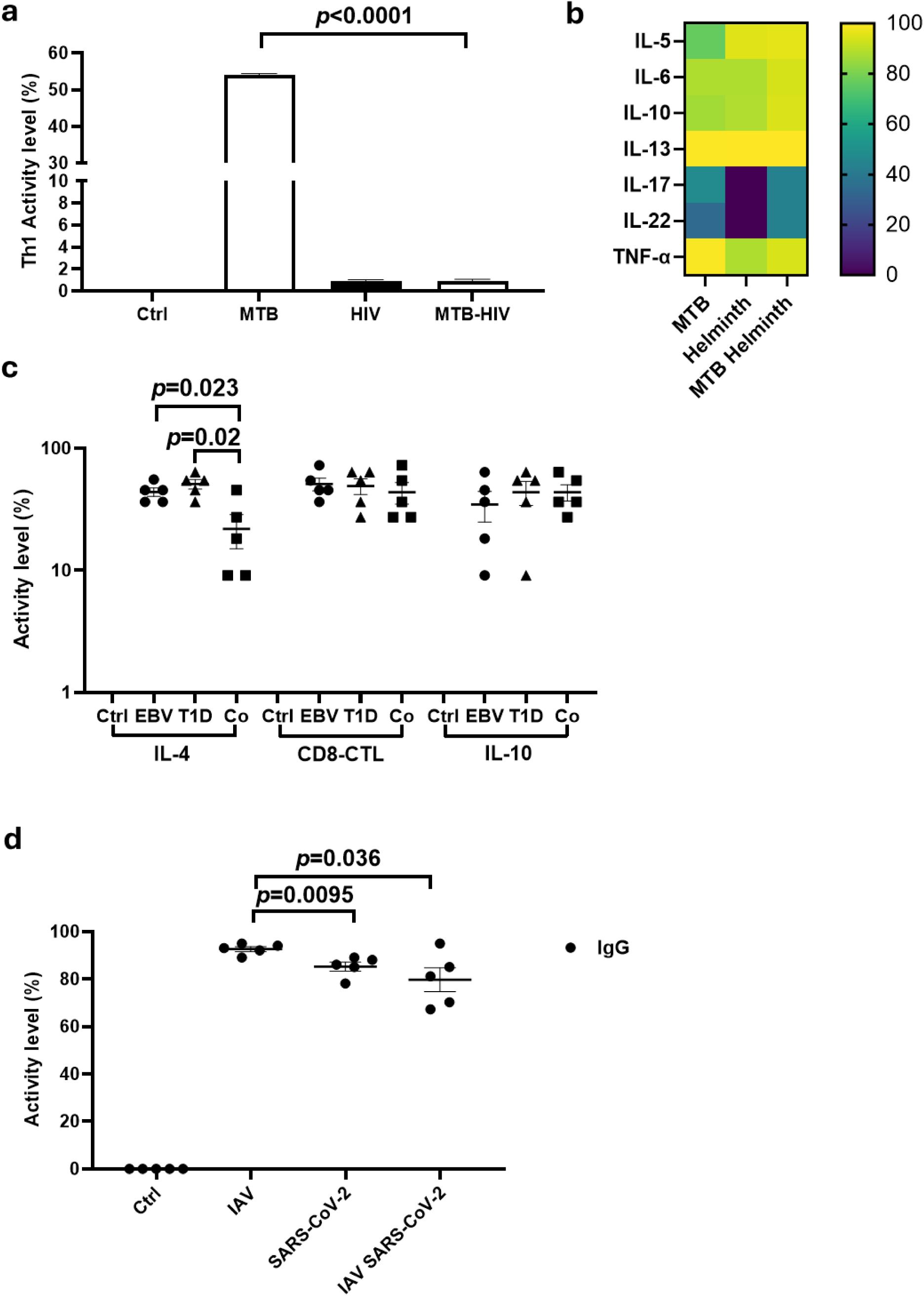
Immune response to coinfections. **a** Activation of CD4+ Th1 in response to MTB, HIV, and MTB-HIV coinfection (replicates n=100 simulations). **b** Differential cytokine response to MTB, Helminth, and MTB-Helminth coinfection. **c** Activity level of IL-4, CD8+ T cells, and IL-10 in response to EBV (circle), T1D (triangle), and coinfection (Co, square) (replicates n=5). **d** Activity level of IgG in response to IAV, SARS-CoV-2, and coinfection. (replicates n=5). All data are presented as mean±SEM, *p*-value determined by unpaired two-tailed t-test.

Additionally, we assessed cytokine responses in MTB-Helminth coinfection. Studies showed that the synergistic effect of MTB-Helminth induces the anti-inflammatory IL-10 along with robust pro-inflammatory responses including IL-5, IL-6, IL-13, IL-17, and IL-22^60,61^. Kathamuthu et al. showed that coinfected patients display an increase of IL-5, IL-13, IL-17, IL-22, and IL-10 compared to individual MTB-infected patients^60^. Our dose-response simulation aligned well with these data, except IL-13 where the simulation failed to demonstrate any difference in activity levels between the single and coinfection conditions (Fig. 4b). In another study, Bewket et al. showed that IL-6 and TNF-α are higher in MTB single infection and coinfection regardless of Helminth infection^61^. Again, our simulation properly predicted the increase of IL-6 and TNF-α in MTB-Helminth coinfection (Fig. 4b).

In the third experiment, we evaluated EBV infection in the setting of T1D using real-time simulations. Klatka et al. showed a decrease in CD8+ T cells and IL-4 secretion in patients with T1D coinfected with EBV, while IL-10 increases in the same coinfected patients^62^. Our model validates the significance of IL-4 decrease when comparing T1D with T1D-EBV coinfected condition (SEM (-22.70± 8.155), p=0.02) or comparing EBV to T1D-EBV coinfection (SEM (-18.35± 6.838), p=0.023); however, our model showed a partial agreement with the data, in which CD8+ T cells and IL-10 levels exhibit an increase without significance (Fig. 4c). The discrepancy between the model simulation and literature might be due to the nature of logical models to be qualitative and non-quantitative.

Finally, we assessed real-time simulations for the IgG responses in cases of both IAV and SARS-CoV-2 coinfection. As demonstrated by Kim et al., IAV exhibited higher IgG titers compared to SARS-CoV-2 single infection and coinfection ^63^. Additionally, there was evidence of IgG impairment *in vivo* in response to coinfection ^63^. Our model corroborated these findings by revealing a significant reduction in IgG levels when comparing IAV to coinfection (SEM (-12.9 ± 5.14), p=0.036), as well as IAV to SARS-CoV-2 alone (SEM (-7.36 ± 2.169), p=0.0095). IgG in IAV-SARS-CoV-2 coinfection is lower than SARS-CoV-2 single infection; however, we did not observe a significant difference in activity level (Fig. 4d).

These studies provide evidence that the model can mostly replicate the immune response during coinfections. Importantly, the model correctly simulates the activity of various cells, cytokines, and Igs in coinfection, which aligns with the literature. These findings pose important solutions for addressing questions related to coinfection.

### Case study 2: Immune response in LTx associated with CMV, EBV, and SARS-CoV-2

LTx presents a complex scenario wherein recipients face the challenge of balancing immune suppression to prevent graft rejection with the need to mount effective immune responses against infectious agents, specifically CMV and EBV. Lung transplant recipients are particularly susceptible to CMV reactivation due to the intense immunosuppressive regimens required to prevent allograft rejection^64^. Similarly, EBV infections can result in severe outcomes in post-transplant patients^65^. SARS-CoV-2 infections’ effects on the respiratory system pose a grave threat to those with compromised lung function, increasing the concerns about the susceptibility and outcomes of lung transplant recipients if infected^66^. Indeed, the COVID-19 pandemic presented unprecedented challenges for lung transplant recipients as shown in clinical studies where lung transplant patients infected with SARS-CoV-2 had higher mortality rates and the survivor exhibited decreased lung function even after recovery^67,68^. The delicate balance between immune response to prevent rejection and immune activation to combat infections like CMV, EBV, and SARS-CoV-2 underscores the importance of tailored management strategies and vigilant monitoring in lung transplant recipients^69^.

Here, we conducted a comparative *in silico* simulation of LTx conditions under three distinct viruses: CMV, EBV, and SARS-CoV-2. Previous studies have established that CD8+ T cells play dual roles, promoting rejection in LTx^70^ while also controlling viremia, primarily through their cytotoxic activity^71,72^. Zaffiri et al. demonstrated that CD8+ T cell levels are higher in LTx without EBV infection compared to those with EBV^73^. Others have published that the frequency of CD8+ T cells remained stable over time in LTx patients regardless of CMV infection^74^. Interestingly, in SARS-CoV-2 infection, CD8+ T cell levels decrease following vaccination^75^. Due to the lack of CD8+ T cell experimental validations in dual conditions of LTx and SARS-CoV-2, we validated our findings based on SARS-CoV-2 vaccination data. Our simulation revealed that the CD8+ T cell response in LTx and LTx-CMV conditions was similar (SEM (-0.1120 ± 0.5835), p=0.84), while in LTx-EBV it was lower compared to LTx alone (SEM (-48.31 ± 1.735), p<0.0001), which is consistent with previous findings. Notably, in the SARS-CoV-2 condition, the CD8+ T cell response decreased compared to LTx alone (SEM (-44.62 ± 1.769), p<0.0001), mirroring trends observed in vaccination data (Fig. 5a).

**Figure 5:**
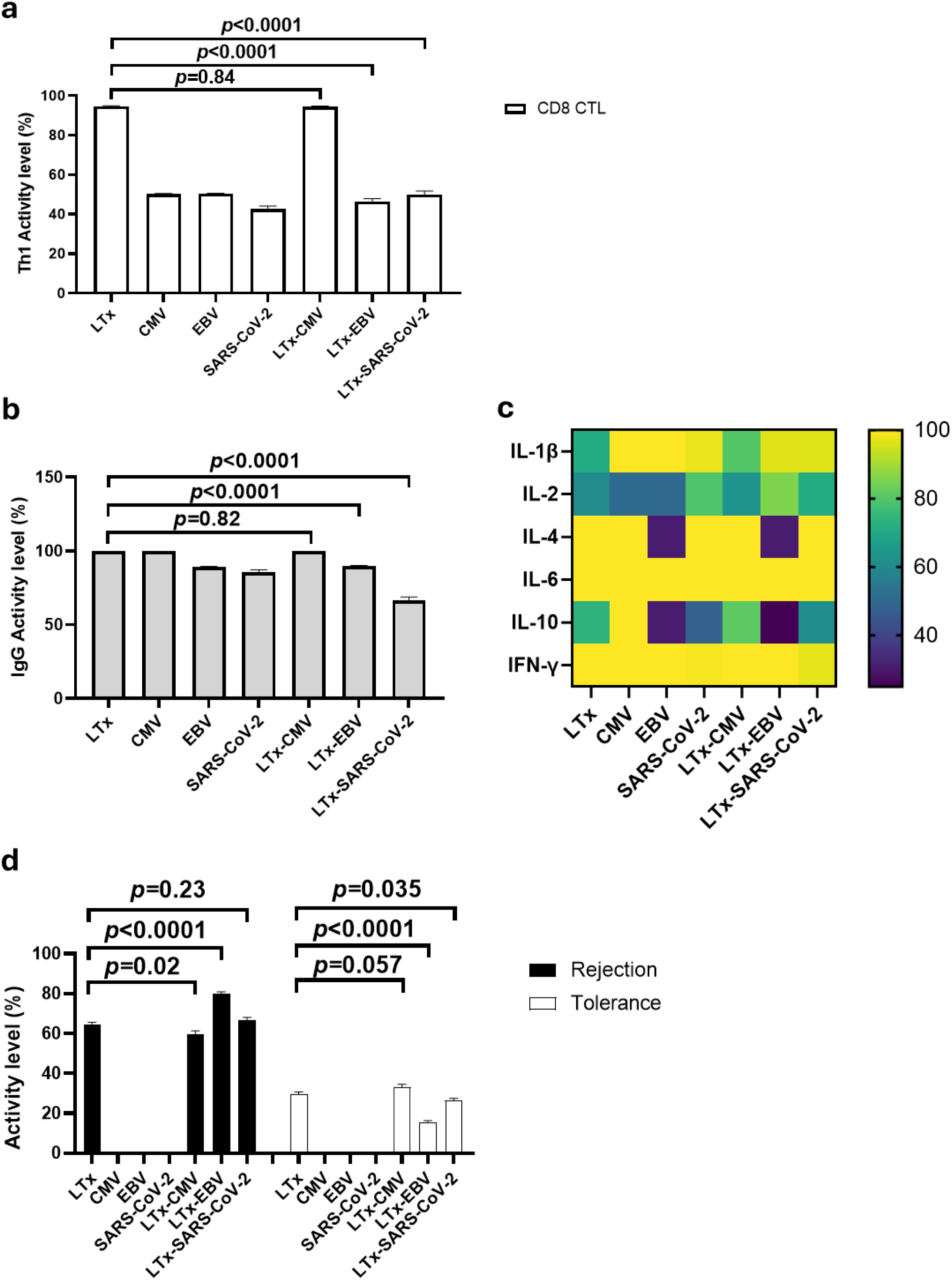
Immune response to LTx and CMV, EBV, and SARS-CoV-2 infections using dose-response analysis. **a** CD8+ T cell response to various single and dual infections. **b** Activity levels of IgG. **c** Cytokine profile of immune response to various pathogens and combinatorial disease conditions. **d** Rejection and tolerance phenotype response in single and dual conditions. The activity levels represent the average value of 100 simulation results triggered by each disease condition. Simulations were performed at 67-100% pathogen activity level. Data are presented as mean±SEM, *p*-value determined by unpaired two-tailed t-test.

Next, we assessed the IgG responses under comparable environmental conditions since it is generally used as a quantitative biomarker of pathogen infection. Similar to the dual nature of CD8+ T cells in LTx, the IgG protects against infections, while also contributing to rejection by targeting donor-specific antigens^76^. LTx patients demonstrated robust IgG responses against CMV, targeting a range of epitopes, with the response correlating with viral load^77^. Conversely, in EBV infection, patients exhibited limited IgG responses, primarily attributable to the target cells of EBV being B cells, which can differentiate into plasma cells. This observation is consistent with our model simulations (SEM (-10.31 ± 0.4797), p<0.0001)^78^. Additionally, several cohorts of lung transplant recipients receiving SARS-CoV-2 vaccines failed to mount a sufficient IgG response ^79^. Our findings complement the published literature showing that the IgG drop is mostly observed in SARS-CoV-2 infection (SEM (-33.26 ± 2.032), p<0.0001) (Fig. 5b).

We next assessed different cytokine responses for their pivotal role in promoting inflammation and tolerance^80^. As shown in Fig 3, the pro-inflammatory IL-1ß, IL-6, and IFN-γ are highly represented across all pathogens. Notably, in LTx, these cytokines have been implicated in post-transplant complications^81–83^. Our simulation results (Fig. 5c) aligned with clinical observations for individual conditions (LTx, CMV, EBV, and SARS-CoV-2). In dual LTx-CMV, LTx-EBV, and LTx-SARS-CoV-2 conditions, IL-6 and IFN-γ mirrored the patterns observed in single conditions, except for IL-1ß. The activity level of IL-1ß in LTx-CMV behaves like the LTx profile and not CMV single infection. Additionally, a subtle reduction in IL-1ß was evident in LTx-EBV compared to EBV alone.

Whitehead et al. observed an elevation of IL-4 levels in bronchoalveolar lavage samples from patients experiencing acute allograft rejection compared to those without rejection^84^. In our simulation, IL-4 was active in all conditions except for those associated with EBV infection, where its presence was notably reduced (31% for both EBV alone and LTx-EBV), suggesting a potential dependency of IL-4 secretion on EBV infection. This observation is consistent with prior research by Buidiani et al., which demonstrated decreased IL-4 expression in EBV-associated infections^42^.

Numerous studies have highlighted the presence of IL-10 and IL-2 in the majority of LTx patient samples, regardless of the presence or absence of complications, with occasional instances of low expression^81,83–86^. Our predictive analysis revealed a diminished expression of IL-10 and IL-2 across all conditions, except for IL-10 in CMV (Fig. 5c). Of note, IL-10 was previously shown to increase during CMV infection^87^; however, our simulation indicated that CMV infection does not significantly impact IL-10 levels in LTx coinfection.

To assess the clinical impact of the immune response in LTx, we incorporated two key phenotypes into the LTx environment: rejection and tolerance. Our simulation verified that both rejection and tolerance components are inactive in single infections (CMV, EBV, and SARS-CoV-2) (Fig. 5d). Furthermore, our *in silico* model predicted that EBV infection elevates the likelihood of rejection (SEM (15.39 ± 1.691), p<0.0001), while SARS-CoV-2 doesn’t show a significant effect (SEM (2.252 ± 1.871), p=0.23). In contrast, CMV infection decreases the likelihood of rejection (SEM (-4.774 ± 2.029), p=0.02). However, the tolerance phenotype is poor when associated with SARS-CoV-2 (SEM (-3.212 ± 1.517), p=0.035) and does not improve with CMV (SEM (3.466 ± 1.812), p=0.057). Our model highlights the heterogeneous responses during viral infection and confirms the negative impact of CMV^88^, EBV^89^, and SARS-CoV-2^90^ infection in LTx patients. Notably, to date, no clinical study has directly compared the outcomes of LTx patients infected with these pathogens, and our model may help to bridge these gaps.

In summary, our model demonstrated the capability to interrogate the immune digital twin within a clinical context, elucidating immune cell behavior in response to multiple infections simultaneously.

## Discussion

The immune system is a highly complex and dynamic network, spanning multiple temporal scales and levels of biological organization. Developing an accurate and comprehensive digital twin of the immune system necessitates a solid foundation that incorporates the essential cellular components and their interactions. The logical model and blueprint presented in this study serve as this foundation, ensuring that the innate and adaptive immune cells, cytokines, immunoglobulins, and other components of the immune system are accurately represented before introducing additional levels of complexity. As the community continues its efforts towards implementing immune digital twins and including other levels of biological organization, the blueprint will serve as a guide for constructing the multi-scale model. The immune system is an area of active research stemming from many different disciplines, including immunology and computational biology. A comprehensive blueprint of the immune system can facilitate collaboration among these researchers by providing a common framework and language for understanding the system’s complexity and interactions.

We validate the cellular-level logical model against available experimental data to assess their accuracy in representing the immune system’s response to different modeled pathogens, which include bacteria, viruses, and parasites. This assessment allowed us to gauge how well the model captures the complexities and interactions within the immune system when faced with different types of infections. For instance, the model was able to reproduce the activation and differentiation of T cells in response to viral infections, as well as the recruitment and activation of neutrophils during bacterial infections.

Despite the compelling advantages of logical modeling, there are some limitations to this approach. For example, logical models are qualitative and therefore lack the quantitative precision required to model the kinetics of immune responses. Furthermore, the construction of logical models is heavily dependent on the current state of knowledge and may lack components or interactions that have not yet been discovered or fully understood. On the other hand, logical models provide numerous benefits, particularly in their ability to capture complex systems in a simplified and computationally efficient manner. They are especially useful for systems with incomplete kinetic data, as they allow for studying system-level behavior based on the known structure of the regulatory network. Logical models can be used to generate hypotheses about the system’s behavior, which can then be tested experimentally. Moreover, they are valuable tools for identifying key regulatory components and potential intervention points in the system.

While the immune digital blueprint model provides a comprehensive representation of many key immune system components and their interactions, it is important to acknowledge that it is not an exhaustive depiction of the entire immune system. There are still many cellular and molecular components, as well as interactions, that have not been included in the current model. The immune system model is also limited to pathogens as triggers for an immune response. For example, other specialized subsets of T cells, such as follicular helper T cells and resident memory T cells, are not explicitly represented in the current model. Similarly, the model does not include immune cell homing, chemotaxis, and the influence of the local tissue environment on cell behavior. Moreover, some cellular components (e.g., innate lymphoid cells) and molecular factors (such as cytokines) have not been thoroughly investigated experimentally, constraining our biological understanding and the model’s accuracy. Regarding infectious diseases, the focus is on the most documented pathogens, rather than their strains. However, the model is primarily designed to capture the core, broadly applicable aspects of the immune system functioning against a variety of insults. Our validation exercises have shown that these key immune responses can be accurately represented even without these additional cell types and states. Thus, while these components could provide additional granularity and potentially enable the model to address more specific questions in future iterations, their absence does not undermine the model’s current accuracy or utility in simulating the general patterns of the immune response. Furthermore, the model’s accessibility in Cell Collective and through SBML provides an opportunity for the community to continue to refine and expand the model to enable simulations of the immune response to other insults, such as allergies, autoimmune conditions, and trauma.

The current blueprint model focuses primarily on immune cell-level communication, including direct and indirect interactions (e.g., cytokines, growth factors, immunoglobulins). To provide a more comprehensive understanding of the immune system, future work should aim to integrate multi-scale modeling approaches, encompassing genetic, molecular, cellular, tissue, organ, and organism levels. By incorporating information from these various scales, researchers can develop a more holistic understanding of the immune system and its role in health and disease. Including physiologically based pharmacokinetics/pharmacodynamics, for instance, would allow the model to predict the effects of drug interventions on the immune system and how these effects might vary among different individuals or under different physiological conditions.

We recently created the first draft of a multicellular, multi-scale, and multi-approach computational model of CD4+ T cells^91^, which can serve as a computational framework for implementing additional scales atop the cellular-level blueprint model presented here. This multi-scale framework integrates physiological (ordinary differential equations), cellular (agent-based approaches), molecular (stochastic logical approach), and genome-level (constraint-based approach) models with heterogeneous high-throughput datasets and bioinformatics algorithms. The model also represents the target cells of initial insult/infection (e.g., epithelial lung cells for infectious diseases), lymphoid tissues connected to the site of insult (a micro-environment where antigens can stimulate antigens), and the circulatory system. The multi-scale and multicellular model demonstrates mathematically and computationally how information flows within and across scales in a single integrated framework, validated by reproducing observed responses of naive CD4+ T cells to different combinations of cytokines.^91^

The presented cellular-level blueprint immune system model will guide the expansion of the aforementioned multi-scale, multicellular framework by expanding it and incorporating additional immune cell-specific sub-models of genome-scale metabolism and signal transduction. While a few such models already exist (e.g., signal transduction network models of antigen-presenting cells^30^, CD4+ effector T cells^92^, and macrophages^93^ and constraint-based metabolic models of Th1, Th2, Th17, and regulatory T cells ^72,94^ and macrophages^95^), the majority of sub-models of other immune cells will need to be developed.

Future multi-scale general-purpose immune digital twins can be applied to many immune-related conditions, including autoimmune diseases and primary immune disorders, infectious diseases, cancer, immunotherapy, chronic diseases, wound healing, transplantation, and trauma responses. By considering the immune digital twin scope within these areas, researchers can explore the immune system’s involvement in these processes and identify potential interventions to improve patient outcomes. For example, the model could be used to simulate the immune response during wound healing to optimize wound care strategies, or to predict the risk of graft rejection in transplantation settings, leading to better patient management and improved clinical outcomes.

## Conclusion

This study provides a comprehensive and simulatable logical model of the immune system, serving as the first blueprint for an immune system digital twin. This blueprint integrates 51 innate and adaptive immune cells, 37 secretory factors, and 11 different disease conditions, providing a solid foundation for developing a multi-scale model. We demonstrated the model’s potential in characterizing system-wide immune responses to various disease conditions. By making the model available in easy-to-use formats directly in the Cell Collective platform, it can be easily further expanded by the community.

The presented cellular-level blueprint of the immune system represents a significant step toward the development of general-purpose immune digital twins. The development and application of immune digital twins have far-reaching implications for the future of digital twin technology in life sciences and healthcare. As digital twins continue to advance, they have the potential to advance patient care and accelerate the transition toward precision medicine. By integrating diverse data sources and providing comprehensive, dynamic models, digital twins enable researchers and healthcare professionals to investigate system behavior, optimize treatments, and develop innovative solutions to pressing medical challenges. Furthermore, digital twins can facilitate collaboration among researchers from different disciplines, providing a common framework and language to understand the complexities and interactions of biological systems.

## Author Contribution Statement

Conceptualization, RA and TH; Methodology and Validation, RA, SSA, LM, and BLP; Resources, LM, DS, KP, and RH; Supervision RA, SSA, BLP, and TH; Writing - Original draft, RA, SSA, BLP, and TH; Writing - Review and Editing, RA, SSA, BLP, and TH; Funding Acquisition, TH. All authors read and approved the final manuscript.

## Acknowledgements

This study was funded by an NIH Grant #R35GM119770 and a University of Nebraska-Lincoln Grand Challenges Catalyst Award to Dr. Tomas Helikar. We thank our scientific writer, Dr. Lindsey Kennedy (University of Nebraska-Lincoln), for editing and formatting the manuscript.

## Data Availability

The mechanistic model is available on the Cell Collective platform at request.

